# Resolving Filament Level Mechanics in Collagen Networks using Activity Microscopy

**DOI:** 10.1101/382903

**Authors:** Emanuel N. Lissek, Tobias F. Bartsch, Ernst-Ludwig Florin

## Abstract

Collagen is the most abundant protein in humans and the primary component of the extracellular matrix, a meshwork of biopolymer networks, which provides structure and integrity to tissues. Its mechanical properties profoundly influence the fate of cells. The cell-matrix interaction, however, is not well understood due to a lack of experimental techniques to study the mechanical interplay between cells and their local environment. Here we introduce Activity Microscopy, a new way to visualize local network mechanics with single filament resolution. Using collagen I networks *in vitro*, we localize fibril positions in two-dimensional slices through the network with nanometer precision and quantify the fibrils’ transverse thermal fluctuations with megahertz bandwidth. Using a fibril’s thermal fluctuations as an indicator for its tension, we find a heterogeneous stress distribution, where “cold” fibrils with small thermal fluctuations surround regions of highly fluctuating “hot” fibrils. We seed HeLa cells into collagen networks and quantify the anisotropy in the propagation of their forces.

## Introduction

Filamentous biopolymer networks fulfill a plethora of mechanical functions both inside and outside of cells. Intracellular networks impart motility and mechanical strength to cells^1^, while the networks of the extracellular matrix (ECM) provide integrity to tissues and entire organs^2^. A particular biopolymer network can fulfill many distinct functions, often achieved by the same basic building blocks arranged in different architectures. For example, collagen, the most prominent component of the extracellular matrix, can be arranged into networks of fibers with wildly diverse densities and connectedness, from the loose and soft elastic networks that form the interstitial matrix of the skin to densely packed mineralized fibers in bones^3^. Many physiological processes are regulated by the stiffness of the ECM, such as cellular migration, differentiation or proliferation^4-6^. The stiffness also plays a role in the progression of skin cancer, as cancerous cells remodel the collagen network of the extracellular matrix to reach blood vessels^7^.

Filamentous biopolymer networks are used by engineers as scaffolds to build artificial tissues that mimic true physiological mechanical properties. However, such approaches remain challenging without a better understanding of the complex interplay between individual filament properties, network architecture and mechanical function^8,9^. Since the typical pore size of these networks is on the same order as the size of cells embedded in the network, cells interact mainly with the individual filaments that surround them, rather than with the global network. To migrate through the extracellular matrix, for example, cells must either squeeze through pores in the network or remodel fibers. Systematic progress in understanding such local cell-matrix interaction is hindered by a lack of techniques that can simultaneously resolve the local network architecture and the interaction forces between cells and filaments.

Currently, the three-dimensional architecture of networks and their interaction with cells can be resolved by confocal microscopy, either in fluorescence or reflection mode, or by light sheet microscopy^10^. These imaging modes do not, however, directly measure the mechanics of the interaction. Forces generated by cells can be indirectly quantified from network deformation data by traction-force microscopy (TFM), provided that the mechanical properties of the filaments and their connectedness are known. Recently, Steinwachs *et al*.^11^ computed for the first time^12^ the forces that breast carcinoma cells apply to biopolymer networks designed to mimic the cells’ physiological surroundings. Forces were calculated from network deformations around the cell using a finite-element approach based on a constitutive equation that captures the complexity of the surrounding network. A drawback of this method is that the measurement of network deformations requires knowledge of the force free state, which is achieved by disassembling the force generating filamentous actin network within the cells. The method then assumes that all deformations are elastic, which neglects plastic deformations that have been observed to occur in the ECM^13,14^. There currently exists no method capable of continuously quantifying cell-matrix interactions with single filament resolution and without the assumption of elastic deformations.

Here we introduce Activity Microscopy, a method for measuring the precise location and the lateral thermal fluctuations of filaments in fibrous biopolymer networks with subnanometer precision and megahertz bandwidth. The magnitude of the lateral fluctuations is a function of filament tension, bending stiffness, network architecture, and, possibly, fluctuating active forces. Therefore, lateral fluctuations are a direct measure for cell-matrix interactions. For an *in vitro* collagen I network, we localize fibrils with nanometer precision and measure their fluctuations along their contour. We find that tension bearing fibrils with small fluctuations surround pockets of weakly loaded fibrils with larger fluctuations. To demonstrate the sensitivity of Activity Microscopy to changes in fibril stiffness, we observe the reduction of fibril fluctuations after cross-linking with glutaraldehyde. Finally, we seed collagen matrices with HeLa cells and measure the reduction in magnitude and the increase in asymmetry of fluctuations near cells.

## Results

### Principle of Activity Microscopy

Activity microscopy aims to visualize the contribution of every filament to a network’s macroscopic mechanical response while simultaneously providing precise information about the network architecture. The filament bending stiffness, stretching and compression forces, and network connectivity affect transverse filament fluctuations, which can either be thermally driven or caused by active elements in the network. We recently demonstrated that thermally driven filament fluctuations can be quantified even in “athermal” collagen I networks^15^ for which transverse fluctuations are not expected to contribute to the mechanical response^16^. Here, we develop this method further and introduce two-dimensional Activity Microscopy. We image areas that include many pore diameters and thus provide insight into how individual filaments contribute to the network’s overall mechanical response.

For Activity Microscopy imaging, a near infrared laser beam is focused into a sample chamber filled with a filament network. The forward scattered and unscattered laser light is then collected and guided onto a quadrant photo diode (QPD) (**Fig. 1a**). This detection scheme is commonly used in optical trapping experiments^17,18^. The QPD outputs two signals, which are calculated from the voltage differences between the quadrants, *S_x_* = (*S_I_* + *S_III_*) – (*S_II_* + *S_IV_*) and *S_y_* = (*S_I_* + *S_II_*) – (*S_III_* + *S_IV_*) (**Fig. 1a,b**). In optical trapping experiments, *S_x_* and *S_y_* can be directly related to the *x* – and *y*-position of a trapped particle. However, in the case of filaments, the signals are not independent but depend on the orientation of the filament relative to the QPD (**Supplementary Fig. 1**).

**Figure 1.**
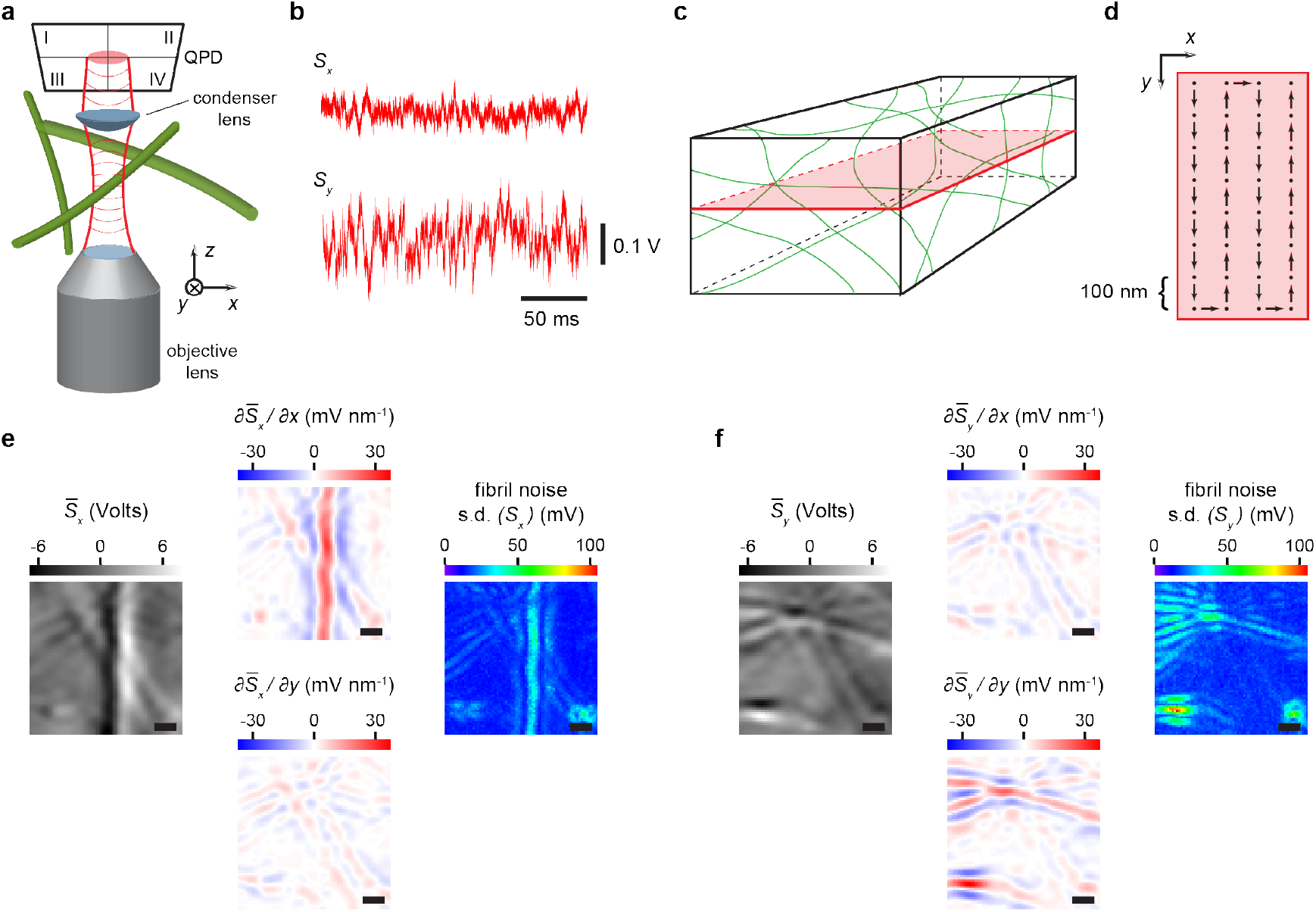
Activity Microscopy. (**a**) A near infrared laser beam is focused into the sample filled with a collagen I *in vitro* network (green: fibrils). The forward scattered light from a fibril is collected by a condenser lens and guided onto a quadrant-photo diode (QPD) where it interferes with the unscattered portion of the laser beam. A time series of lateral voltage signals *S_x_* = (*S_I_* + *S_III_*) − (*S_II_* + *S_IV_*) and *S_y_* = (*S_I_* + *S_II_*) − (*S_III_* + *S_IV_*), recorded at one position are shown in **b**. (**c, d**) For imaging, a plane of interest in the sample is selected and scanned line by line while recording time series at each position (black dots). The mean detector signal *S̄_x_* is shown in the left panel of **e**. To find the fibrils’ axes, the spatial derivatives *∂S̄_x_*/*∂x* and *∂S̄_x_*/*∂y* are calculated, termed detector sensitivity (middle panel). The raw transverse fibril noise is calculated from the time series of *S_x_* and displayed as the standard deviation (s.d.) (right panel). **f** shows the averaged detector signal, derivatives and transverse fibril noise for the *y*-signal, which has its maximal sensitivity for fibrils oriented along the horizontal axis. Scale bars in **e** and **f** are 1 μ*m*. The collagen network was polymerized according to protocol I (see Methods). Distance between scanning lines and dots in **d** is 100 *nm*. Time series at each location in **e** and **f** is 100 *ms* long.

For two-dimensional imaging, a plane of interest is selected (**Fig. 1c**) and raster scanned by translating the sample in steps (**Fig. 1d**), which are chosen to be smaller than the diffraction limited spot diameter. **Figure 1e** and **f** show the average detector signals, the calculated detector sensitivities and the raw noise signal for the example of an *in vitro* collagen I network. A total area of 7 μ*m* × 7 μ*m* was scanned with a step width of 100 *nm*. At each point, a time series of 100 *ms* was recorded at 100 *kS s*^−1^. The left panel in **Figure 1e** shows the averaged detector signal *S̄_x_* along the *x*-axis from which the detector sensitivities *∂S̄_x_*/*∂x* and *∂S̄_x_*/*∂y* (mid panels) are calculated. Clearly, *S_x_* is mainly sensitive to fibrils that are oriented vertically. The highest sensitivity is measured when the laser beam is centered on the fibril (broad vertical line). This is expected from the detector response to a fibril, which has its maximal slope at the midpoint of the fibril (**Supplementary Fig. 1**). Thus, the location of maximal sensitivity can in turn be used to determine the precise position of the fibril axis. The fibril’s transverse position fluctuations can be calculated from the standard deviation s.d. of the times series of *S_x_* and *S_y_*. Since the highest detector sensitivity leads to the largest raw position noise for a given filament fluctuation amplitude, the maxima in the raw noise image (right panel) also indicate the positions along the fibril axis.

The fibril noise images in **Figure 1e** and **f** show “ghost filaments” that run next to the main fibril axis. They result from the characteristic detector response of a fibril. Besides the maximal detector sensitivity when the fibril is directly in focus, the detector also has significant sensitivities to the left and right of the center (**Supplementary Fig. 1c**, dashed lines). These regions of the detector response with negative slope are clearly visible in the detector sensitivity panels as blue lines, i.e. negative sensitivities, that follow the fibril on either side. Since negative sensitivities can be easily identified, this detection artifact can be corrected for by removing signals originating from locations in the sample with negative sensitivities (**Supplementary Fig. 2**).

### Finding Filament Axes and Orientations

Point-by-point Activity Microscopy imaging is time consuming because at each point in the region of interest a time series must be recorded for a sufficiently long time to characterize the local fluctuations of the filaments. Given an approximate pore size of 2 – 3 μ*m* for the collagen concentration used here (2.4 *mg ml*^−1^) (ref. 19), most of the scanning time is spent inside pores. Additionally, fibril fluctuation amplitudes can only change along the contour of a filament, but not transversally to the contour. Thus, data from the same position along the contour of the filament, but at different positions transversal to its axis, yield the same fluctuation amplitude as long as the filament remains within the linear range of the detector (**Supplementary Fig. 3**). For characterizing the fluctuation of each fibril in the network, it is therefore sufficient to measure fibril fluctuations only at points along the fibril axis. To implement such an imaging algorithm, the locations and orientations of all fibrils in a network must be known. We achieve this by rapidly pre-scanning the network (“fast scan”), only recording time series sufficiently long to determine the average detector signals *S̄_x_* and *S̄_y_*.

**Figure 2a** and **c** show the results for a time series with a length of 5 *ms* (**Supplementary Note 1**). There is no obvious degradation in the *S̄_x_* and *S̄_y_* images in comparison to the images shown in **Figure 1e** and **f**. The line profiles of the average detector signals are still smooth (**Fig. 2b**) and the calculated detector sensitivities are essentially noise free (**Fig. 2d,e**). Since the positions of maximal detector sensitivities colocalize with the fibrils’ locations, we can use these images to localize the fibrils. To achieve this, horizontal and vertical line profiles of the *S̄_x_* and *S̄_y_* signals, respectively, are extracted and the locations of the maxima determined. In this way, the fibrils’ axes are found with nanometer precision (**Supplementary Note 1**). **Figure 2f** shows the locations of the fibrils identified using this algorithm on a grid with a pixel size of 100 *nm* × 100 *nm*. Ghost filaments are excluded because they correspond to minima in the detector sensitivity. The orientation of filaments relative to the detector orientation is then determined (see Methods) and displayed color-coded for each filament position. This algorithm reliably finds fibril axes locations and orientation in a 5 μ*m* × 5 μ*m* image in approximately two minutes.

**Figure 2.**
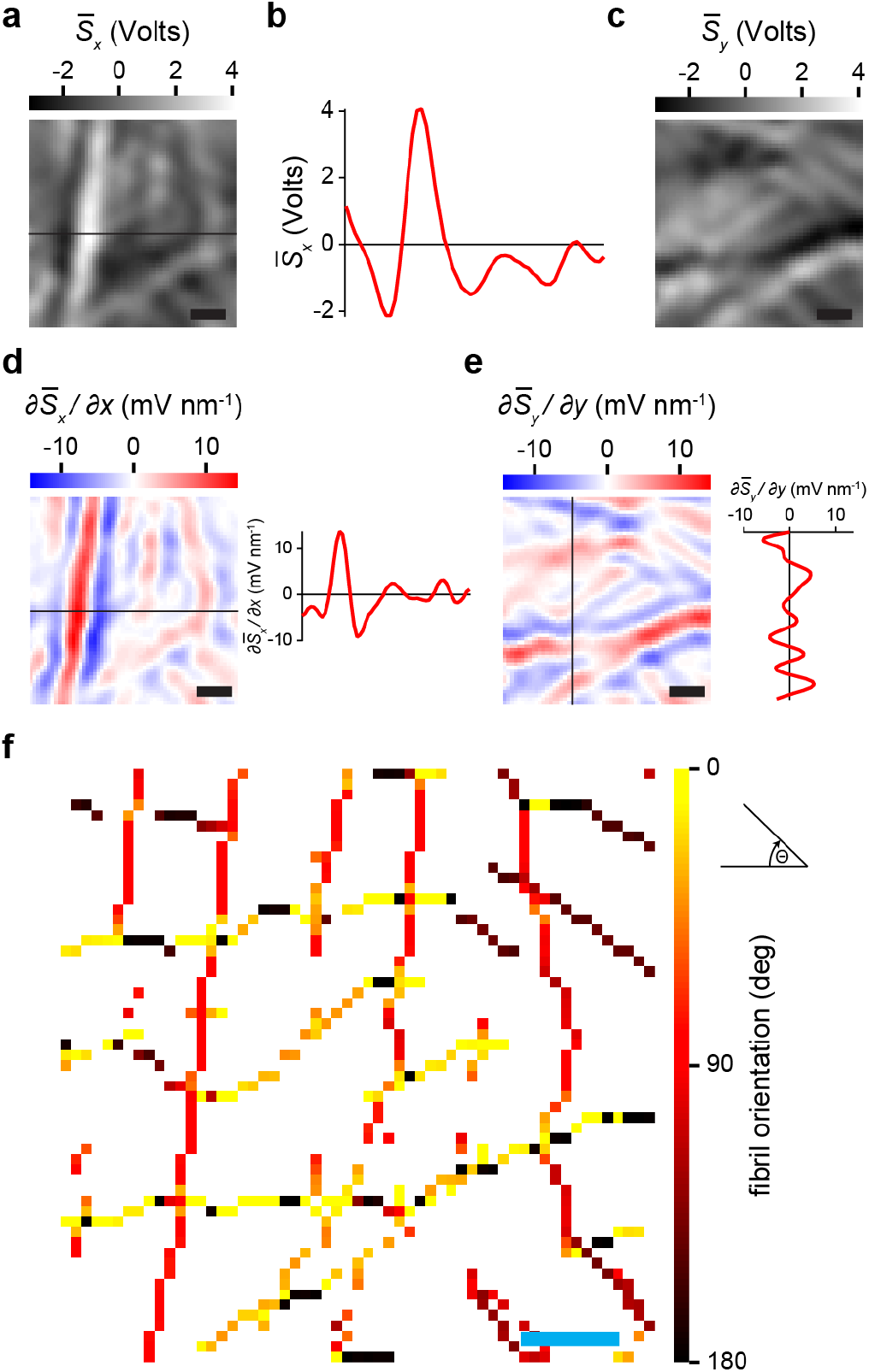
Localization of fibrils. Fibrils are located using the maximum sensitivity values either along the *x* – or *y* – direction depending on their orientation relative to the detector coordinate axes. A line profile of the mean signal *S̄_x_* (black line in **a**) reveals the detector response to an individual fibril (**b**). (**c**) The mean signal *S̄_y_* from the same scan as in a. The detector sensitivities *∂S̄_x_*/*∂x* (**d** and *∂S̄_y_*/*∂y* (*e*) show maxima at the fibrils’ locations (see also representative horizontal and vertical line profiles, respectively). *∂S̄_x_*/*∂x* is used to identify fibrils oriented with an angle ≤ *45* °w.r.t. the vertical direction, while *∂S̄_y_*/*∂y* serves to find fibrils oriented with an angle < 45 ° w.r.t. the horizontal direction. The locations of the fibrils are plotted in **f**. The color indicates the fibril orientation w.r.t. the horizontal direction (see Methods). Scale bars are 1 μ*m*. The collagen network was polymerized according to protocol I (see Methods).

### Long-range imaging

For visualizing how individual filaments contribute to the overall mechanical response of a network, it is necessary to image areas that are large relative to the pore diameter. This is challenging because slow drift of the sample or instrument will lead to increasing discrepancies between the fibril’s positions found by fast scanning, and the locations at which the fluctuations are recorded. To solve this problem, we subdivide the area of interest into smaller tiles of 5 μ*m* × 5 μ*m* and perform Activity Microscopy imaging tile by tile, thus avoiding significant relative drifts. As shown in **Figure 3a**, fibrils can be traced this way over long distances without discontinuities. In each tile, a sequence of four steps is performed. The fast scan with dwell time of 5 *ms* per pixel is recorded and the fibrils’ axes and their orientations are determined (**Fig. 3b**). The list of fibril positions is then used to record fluctuation data for 200 *ms* only directly on the fibrils’ axes. To obtain an accurate detector calibration at the time that fluctuation data are recorded, the detector sensitivity is measured at each pixel of interest by scanning the fibril perpendicularly to its axis through the laser beam (**Fig. 3c,d**). The scan is used to calibrate the raw voltage signals which then provide an accurate measure for the transverse fibril fluctuation at a given position on the filament axis (**Fig. 3e,f**). At every position we chose either the *x-* or *y*-channel, depending on which one has the greater sensitivity. Both channels will pick up the same fluctuation amplitude except for fibrils that are perfectly aligned with one of the detector axes. This selection of channels minimizes the uncertainty in fluctuation measurements (**Supplementary Fig. 4**).

**Figure 3.**
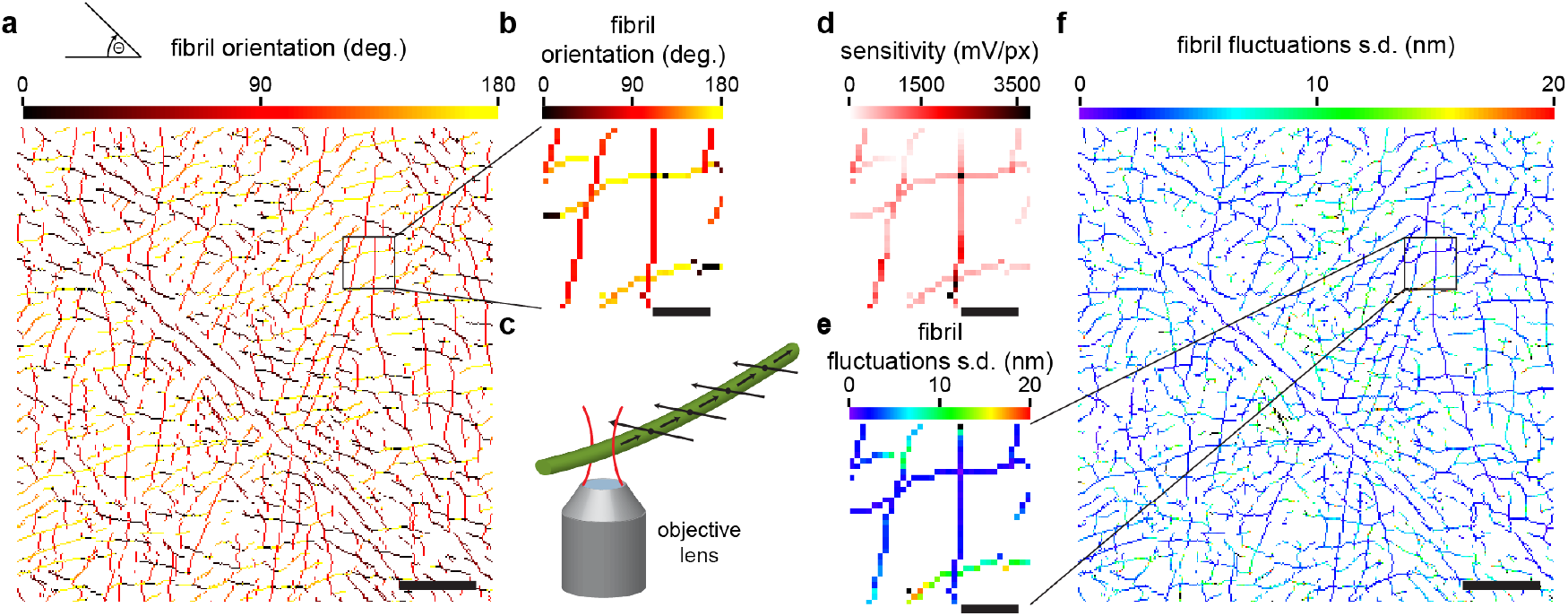
Quantitative imaging of fibril fluctuations. The map of fibril locations and orientations (**a, b**) are used to record fibril fluctuations only at selected points corresponding to fibrils’ axes. In a first step, the detector signal is recalibrated by scanning the laser beam perpendicularly to a fibril axis (**c**). **d** shows the refined detector sensitivities for each fibril position. A 200 *ms* time series is then recorded, and the raw voltage signals are converted into calibrated transverse fibril fluctuations using the measured detector sensitivity. **e** and **f** show the final Activity Microscopy image. Each fibril position is shown color-coded with the magnitude of its transverse fluctuation. Scale bars are 5 μ*m* in **a** and **f**, 1 μ*m* in **b**, **d**, and **e**. The collagen network was polymerized according to protocol I (see Methods).

**Figure 3f** shows a 30 μ*m* × 30 μ*m* Activity Microscopy large scale image of a collagen network, in which the amplitude of the transverse filament fluctuations is color-coded for the range 0 – 20 *nm*. The distribution of fluctuation amplitudes in the network is heterogeneous: a few fibrils with strongly suppressed fluctuations cross the entire network and in-between are pockets of strongly fluctuating fibrils.

### Pore size distribution

Activity Microscopy images of collagen networks suggest a heterogeneous distribution of fibril fluctuations. A few fibrils with strongly suppressed fluctuations divide the field of view, while the regions between them are filled with fibrils with large fluctuations. To quantify this observation, we calculate the average pore size distribution for “hot” (strongly fluctuating) and “cold” (weakly fluctuating) fibrils.

To calculate the pore size distribution, we adapted a robust method for calculating pore sizes from confocal images of collagen networks developed by Mickel *et al*.^19^, which does not rely on specific assumptions or network models. Fibrils are represented by medial axes instead of their diffraction limited image. In our case, fibrils are represented by one-pixel wide lines corresponding to a width of 100 *nm*. For each pixel of the fluid phase, the radius of the largest disk that can be fitted into the pore while still including the pixel, is determined. For pixels within the center of a pore, the maximal disk size is given by the contact of the disk with the medial axes that form the pore. Pixels confined by fibrils in corners will have a smaller radius. The pore size distribution for a network can be summarized by a histogram of radii for all pixels of the fluid phase. The goal in our case is to show that cold and hot filaments have different pore sizes. To achieve this, we divide all fibrils (**Fig. 4a**) into an equal number of cold (blue) and hot (red) filaments based on the cumulative probability of fibril fluctuations (**Fig. 4b**). An equal number of cold and hot fibrils is achieved at a fluctuation threshold of ~ 5 *nm*. A binary image representing the cold and hot fibrils shows again the large-scale pattern of cold filaments surrounding pockets of hot filaments (**Fig. 4c**). To visualize the pore size distribution within an Activity Microscopy image, we plot the maximal pore size, color-coded for every pixel in the fluid phase either for all (**Fig. 4d**), only hot (**Fig. 4e**), or only cold fibrils (**Fig. 4f**). The overall characteristics of the three cases is summarized in the probability density distribution of pore radii (**Fig. 4g**). Except for a few pores, the pore radius for most pores is narrowly distributed around 435 *nm*. Hot filaments alone have a slightly higher peak pore radius of 567 *nm* (**Supplementary Note 2**). In contrast, cold filaments have clearly a much larger average pore radius indicated by the change in color and size of pores (**Fig. 4f**) and the distribution of pore radii with a peak of 897 *nm* (**Fig. 4g**). This confirms quantitatively the observation that cold filaments divide the network into pockets of hot filaments. Assuming that cold filaments are a result of tension along their axes, one could interpret the data as a few filaments carrying tension over long distances, while leaving pockets of relatively low-tension fibrils between them. To our knowledge, such a heterogeneous stress distribution has never been experimentally shown before.

**Figure 4.**
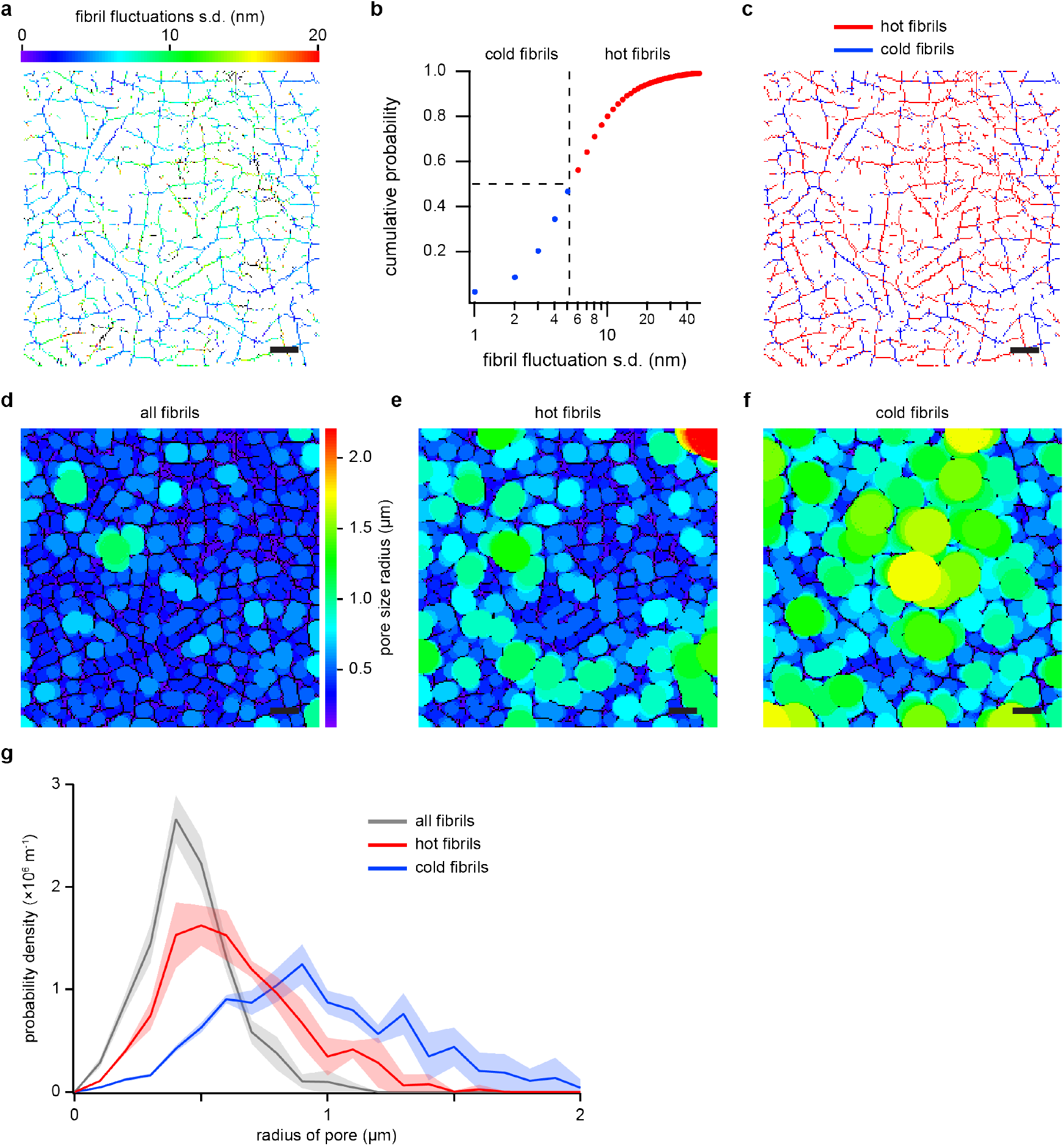
Pore size distribution for strongly fluctuating (hot) versus weakly fluctuating (cold) fibrils. (**a**) Activity Microscopy image (20 μ*m* × 20 μ*m*) of a network of native *in vitro* collagen I fibrils. The cumulative probability of fibril fluctuations for the image shown in a is used to divide fibrils equally into hot (red) and cold (blue) (**b**). The dashed lines indicate the median used to split the fibrils into hot and cold as also displayed in **c**. This image is used to calculate pore sizes between the network’s fibrils (d) using a method described by Mickel *et al*.^19^ (see Methods). The subset of hot fibrils forms smaller pores than the subset of cold fibrils (**e, f**). (**g**) The difference in pore sizes of networks of hot and cold fibrils can be quantified by the probability densities of their pore size radii. Cold fibrils form pores ≈ 1.6 × the size of the pores formed by hot fibrils (shaded region represents s.d. of four independent Activity Microscopy images of two different samples). Scale bars are 2 μ*m*. Collagen was polymerized according to protocol II (see Methods).

### Collagen crosslinking

To demonstrate that Activity Microscopy imaging is sensitive to changes in the mechanics of collagen fibrils, we investigated networks crosslinked with glutaraldehyde, a molecular crosslinking agent commonly used for cell and tissue fixation. Glutaraldehyde crosslinks collagen fibrils internally by coupling individual collagen molecules and connecting the constituent proteins’ surface lysines. It does not, however, change aspects of the network’s structure, such as fiber width and length, or pore size^20^. Therefore, mainly a change in the stiffness of the fibrils and a corresponding reduction in transverse fluctuations are expected. Figure 5a and c show 30 μw*m* × 30 μ*m* Activity Microscopy images of a native collagen network and a crosslinked network, respectively. Both networks were prepared in the same way except for the added crosslinking step (see Methods). The suppression of transverse fibril fluctuation through internal crosslinking of fibrils is immediately visible as a color shift corresponding to smaller fluctuations. To quantify the change in fibril fluctuations, we calculated the probability density distribution from the fluctuation amplitude for all fibrils shown in the Activity Microscopy images. **Figure 5b** shows such a distribution for the native network. The distribution is asymmetric with a maximum around 3 – 4 *nm* and a long tail up to 20 *nm*. After crosslinking, the distribution shows a strong shift towards smaller amplitudes with a maximum around 1 *nm* and a long tail up to 10 *nm* (**Fig. 5d**). These data demonstrate the sensitivity of Activity Microscopy to subtle changes in filament mechanics.

**Figure 5.**
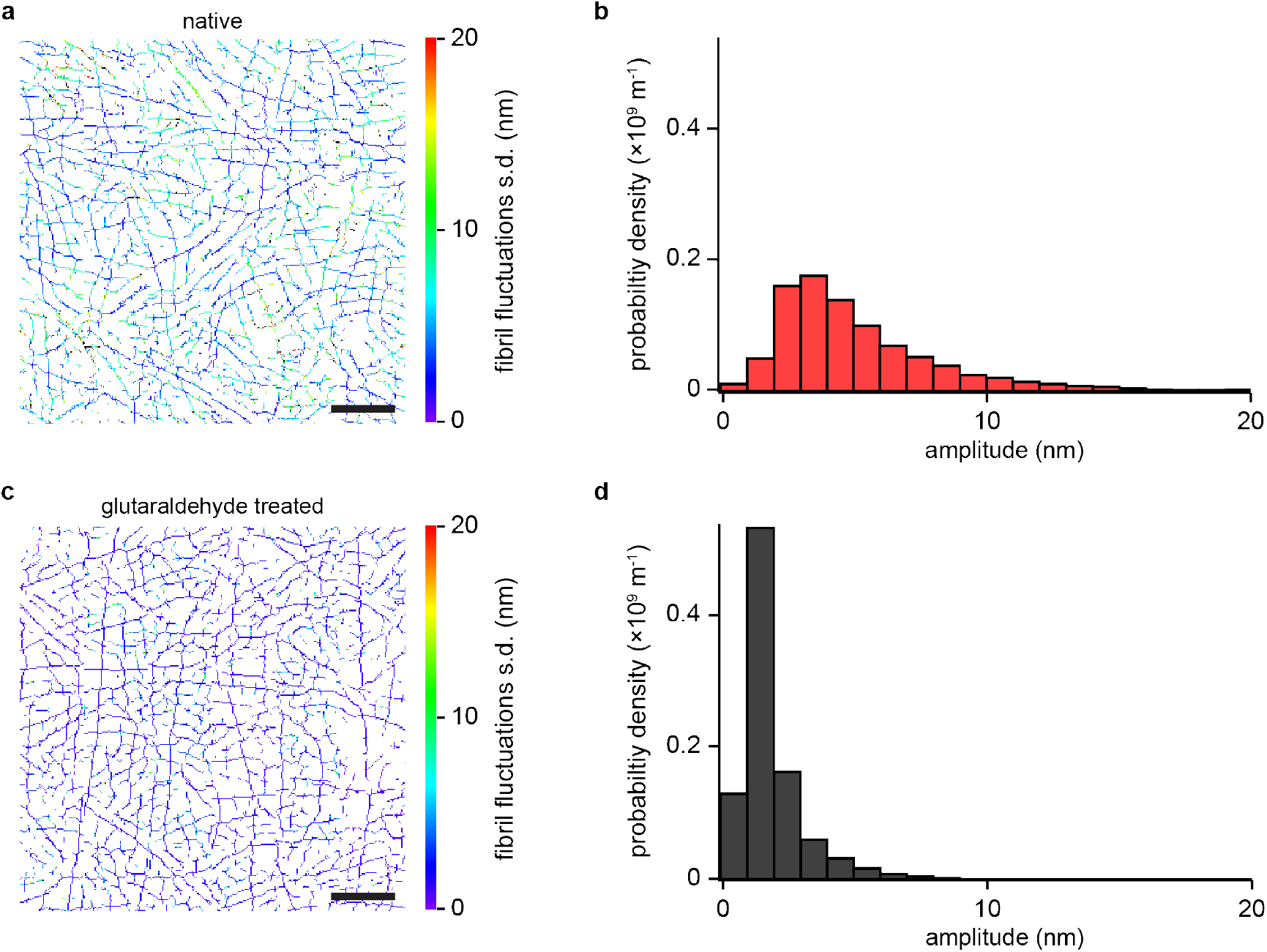
Reduction in collagen fibril fluctuations due to chemical crosslinking. (**a**) Activity Microscopy image of an *in vitro* network of native collagen I fibrils. (**b**) Probability density of fibril fluctuation amplitudes computed from all pixels shown in a. The peak of the distribution is located at 4.7 ± 0.7 *nm* (s.d. of five independent Activity Microscopy images of two different samples). (**c**) Activity Microscopy image of a collagen I network cross-linked with 4 % glutaraldehyde. (**d**) Probability density for the crosslinked network in c. Crosslinking leads to a reduction in fluctuation amplitudes with a new peak location around 1.6 ± 0.4 *nm* (s.d. of 4 independent Activity Microscopy images of 2 different samples). The bin width in **b** and **d** is 1 *nm*. Scale bars are 5 μ*m*. The collagen networks were polymerized according to protocol I (see Methods).

### HeLa cells in a collagen I network: active forces

Cells growing and migrating in collagen matrices apply forces to individual fibrils that result in network deformations and remodeling^4^. To demonstrate that Activity Microscopy is able to quantify and visualize these changes, we seeded HeLa cells into collagen I matrices and measured fibril fluctuations at varying distance from the cells. HeLa cells are known to proliferate but not migrate in collagen^21,22^, yet actively apply forces to the network. We expect fluctuations of fibrils under tension to be reduced, and the tension to fall off with the distance from the cell. **Figure 6a** shows the probability densities of fluctuations amplitudes measured in 10 μ*m* × 10 μ*m* areas next to (black), 50 μ*m* (blue) and 100 μ*m* (red) away from a cell. While there is no significant difference between the distributions measured close to the cell (peak at 3.8 *nm*) and 50 *μm* away (peak at 3.9 *nm*), the distribution shifts towards larger amplitudes at 100 μ*m* away (peak at 4.9 *nm*). Each measurement took ≈ 30 *min* to complete which is sufficiently long to expect changes in the cell’s state. We therefore recorded another distribution next to the cell (yellow). In contrast to the first recording, fluctuations are now strongly suppressed with a peak amplitude of 2.6 *nm* and a corresponding overall contraction of the distribution to smaller amplitudes. To confirm our observation, we repeated the experiments several times and increased the scan area to 20 μ*m* × 20 μ*m* to increase the quality of the probability density distribution. For reference, we measured the distribution for cell free matrices prepared under the same conditions. The average of all measurements for matrices with cells (red) and without cells (black) are shown in **Figure 6b**. The reduction in fluctuation amplitude relative to the cell free matrices confirms that the cells apply tension to the network. This tension seems to fall off within 100 *μm* as the distribution measured 100 μ*m* away from the cell approaches the average distribution measured in cell free matrices. Anisotropy in tension is an additional signature expected for cells applying tension to the surrounding matrix. Cells pulling on the fibrils in their surroundings will deform the network. Fibrils radiating out will carry away and distribute the tension while concentric fibrils will experience reduced tension. To show this anisotropy, we analyzed the fluctuation amplitude dependent on their orientation to the detector coordinate axes. **Figure 6c** shows a polar graph for fluctuations recorded around a point 20 μ*m* away from a cell. Fibrils radiating away from the cell show an ≈ 37 % reduction in fluctuation amplitude relative to fibrils oriented in perpendicular direction. The orientation of the ellipse agrees with the location of the cell relative to the scan area (inset). This can be independently confirmed by light microscopy imaging which allows us to determine the position of the cell relative to the scanned area. As a control, we applied the same analysis algorithm to data recorded on cell free collagen networks. In this case, as expected, fluctuation amplitudes are distributed isotropically (**Fig. 6d**).

**Figure 6.**
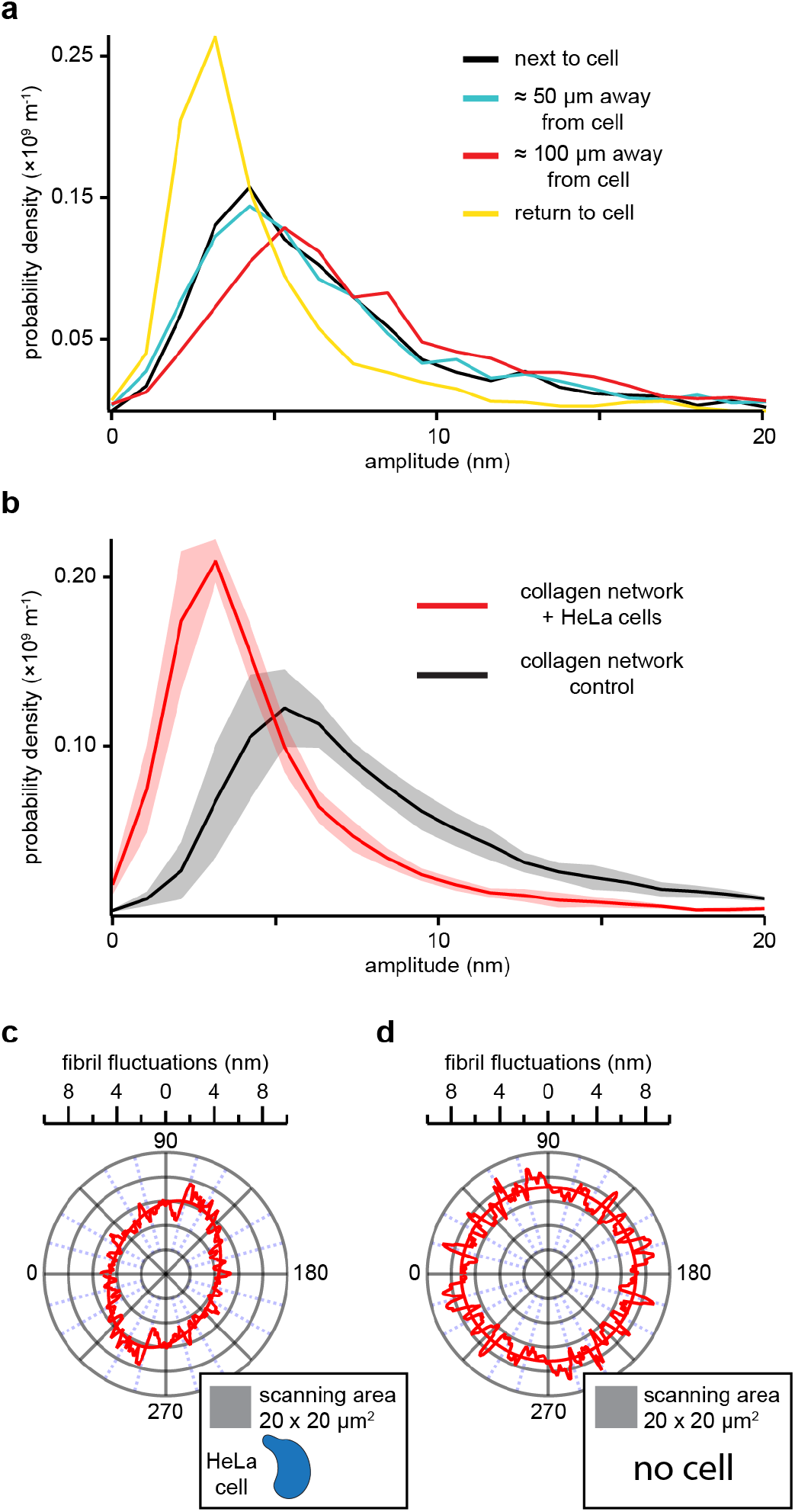
Collagen I network response to embedded HeLa cells. (**a**) Probability density distributions of fibril fluctuation amplitudes at varying distances from a HeLa cell in an *in vitro* collagen I network. Each distribution was computed from a 10 *μm ×* 10 μ*m* Activity Microscopy image. Each scan took ≈ 30 *min* to complete. (**b**) Average probability density distribution (red) calculated from four different 20 μ*m* × 20 μ*m* scans next to HeLa cells (< 5 μ*m*). The shaded region indicates the s.d‥ For reference, an average probability distribution (black) calculated from four scans in a cell free network is shown. The presence of cells shifts the distribution towards smaller fluctuations, from 5.1 *nm* for cell free networks to 2.7 *nm* for networks with embedded HeLa cells. Fibrils analyzed for **b** can be sorted by their angle relative to the detector coordinate axes and the dependence of their fluctuations on this angle can be displayed in a polar graph (**c**). The fluctuation amplitudes reveal that radially oriented fibrils (w.r.t. the cell) display ≈ 37 % weaker fluctuations on average than tangentially oriented fibrils (see inset). No significant anisotropy is observed in cell free networks (**d**). Here, the data from one representative cell free network from **b** were used. All probability densities distributions have a bin width of 1 *nm*. The polar graphs show smoothed raw data (sliding box count = 4) and an elliptical fit. The collagen networks were polymerized according to protocol II (see Methods).

In summary, these data demonstrate that the sensitivity range of Activity Microscopy imaging is well within the range of changes caused by cells to the matrix.

## Discussion

Networks made of stiff biopolymers such as collagen I are usually modeled as athermal^14,23,24^, i.e. the thermal undulations of their constituent filaments are assumed to play no significant role in their mechanical response. Here, we showed that thermally driven transverse filament fluctuations can nevertheless be measured, despite their ångstrøm to nanometer amplitudes. Finding the location of fibrils previous to fluctuation measurements allowed us to reduce two-dimensional imaging to effective one-dimensional fluctuation measurements along a fibril’s contour. In this way, we were able to probe large areas of a network that include many pores, thus bridging the gap between single filament and overall network behavior. The observation of different pore sizes for cold and hot fibrils is one prominent result of this new ability. Further, we calculated probability density distributions of filament fluctuations representing the state of a network and its changes. We demonstrated that these changes can originate from increased stiffness of the individual fibrils in case of crosslinking or from tension applied to the fibrils in case of cells. Thus, Activity Microscopy imaging covers the relevant range of fluctuation amplitudes to characterize networks and their interaction with embedded HeLa cells. In both cases, the thickness of the sample ranged from 0.5 – 1.0 *mm*, sufficiently large to treat the matrix as a three-dimensional network.

Recently, Steinwachs *et al*.^11^ estimated the total force applied by a breast cancer cell to a collagen matrix to be on the order of 50 *nN*. As the pore size of the network is small relative to the cell’s volume, the total force is expected to be distributed over many fibrils, with a maximum force per fibril in the nanonewton range or smaller. To estimate whether transverse fibril fluctuations are still within the range of Activity Microscopy, we calculated the transverse filament fluctuations based on a theory of semiflexible filaments under tension for fibril lengths of 10 – 40 μ*m* and forces up to 10 *nN* (refs. 16,25) (**Supplementary Fig. 5, Supplementary Note 3**). Assuming a detection limit of 1 nm, all fluctuation amplitudes are within the range of Activity Microscopy. Only for the shortest collagen fibrils of length 10 μ*m* and an applied tension of 10 *nN*, the detection threshold of 1 nm is reached. Very small forces do not significantly change the fluctuation amplitudes (see plateaus in **Supplementary Fig. 5**). The minimal detectable tension depends on the fibril length. For example, for collagen fibrils of length *10 μm*, only tensions larger than 100 *pN* significantly change fluctuations amplitudes. For 40 μ*m* fibrils, this threshold is much lower, around tens of piconewtons. These estimates show that Activity Microscopy covers the relevant force range for studying cell-matrix interaction also for cells that are expected to show strong interactions with the surrounding matrix, such as migrating cancer cells^26^.

Imaging speed is another factor required to follow cell-matrix interaction of motile cells. For our study, we chose HeLa cells that are known to interact with collagen matrix, but are unable to migrate^21,22^, leaving sufficient time to image the selected areas of the network around them. Migrating cancer cells, however, move with speeds on the order of micrometers per minute^27^, a time scale on which one would like to monitor their interaction with all fibrils of the matrix. This poses a challenge for Activity Microscopy that can be met with current technology. Pre-scans of volumes around a cell, required for finding all fibrils in a network, can be sped up to less than a minute by using a fast scanning mode of the scanning stage instead of the step by step scanning mode used in this work. This will reduce the precision in localizing the fibrils, but as we pointed out above, the precise knowledge of fibril position is not required for the measurement of fibril fluctuations as long as the fibril remains in the linear range of the detector (**Supplementary Fig. 3**). The characteristic length scale of the network is set by the pore size. The imaging speed can therefore be increased by measuring a fibril’s fluctuations only at single points spaced by the network pore size. Assuming a pore diameter of 2.5 μ*m* (ref. 19), this will reduce the time required to measure fibril fluctuations by a factor of 25, compared to our current stepsize of 100 *nm*. These two improvements in imaging speed might be sufficient for studying the interaction between motile cells and their surrounding matrix.

Activity Microscopy also has broad applications in materials research, where networks are static and hence much easier to image. Collagen networks are extremely stable and therefore large volumes can be imaged. We expect to gain new insight into how single filament mechanics and network architecture lead to macroscopic mechanical properties. This allows for a systematic study of network preparation conditions and a rational design of networks with desired properties. The response to point and shear forces and other network manipulations such as enzymatic activity or chemical treatment by cross-linkers can also be studied.

## Materials and Methods

### High Bandwidth, High Precision Position Detection

Measurements were performed on a custom-built photonic force microscope (PFM) as described in detail by Bartsch *et al*.^15,28^. In brief, the beam of a 1064 *nm* laser (Mephisto, 500 *mW*, Coherent, CA, USA) was expanded and focused through a water immersion objective lens (UPlanSApo, 60 × W, Olympus, Tokyo, Japan) into the sample. The sample chamber was mounted on a three-dimensional nano-positioning stage (Nano-View/M375HS, Mad City Labs, WI, USA), which allowed for the sample to be moved relative to the stationary optical trap. Forward scattered light from a collagen fibril, together with unscattered light of the laser beam was collected by a condenser lens and projected onto a quadrant photodiode (G6849, Hamamatsu Corporation, NJ, USA), where the two waves interfered. The differential signal from the QPD were amplified by custom-built low noise differential amplifiers (SA500, Oeffner MSR, Plankstadt, Germany). This detection scheme has a megahertz electronic bandwidth.

### In vitro Collagen I Networks – Polymerization Protocol I

Collagen networks were prepared and polymerized *in vitro* as described in Bartsch *et al*.^15^. In brief, acid-soluble rat-tail tendon collagen (Collagen I, rat tail, 354236, Corning^®^, NY, USA) and bovine-dermis collagen (Collagen I, bovine, 354231, Corning^®^, NY, USA) were mixed at relative concentrations of 1:2. The mixture was then diluted to a total collagen concentration of 2.4 *mg ml*^−1^ by adding equal parts of 10 × DMEM (D2429, Sigma Aldrich, MO, USA) and 0.27 *M* NaHCO**3**. To induce gel polymerization, the pH of the solution was raised to pH 10 using 1 *M* NaOH. All components were kept on ice during mixing. The mixture was then quickly pipetted into a preassembled sample chamber consisting of a glass coverslip attached by vacuum grease to a metal sample chamber (**Supplementary Fig. 7**) and left to polymerize for ~ 1 *h* at 37° *C* in a humidity-controlled incubator with 5% CO_2_ atmosphere. Polymerizing the network inside the sample chamber ensures its attachment to the coverslip, which is a prerequisite for a mechanically stable assay. Networks were between 500 μ*m* and 1 *mm* thick. After polymerization, the gel was gently rinsed with *1 ml* of *1 ×*Phosphate-buffered saline (PBS). Care was taken to never let the network dry out.

After polymerization of the collagen network, the sample chamber was closed by attachment of a top coverslip (**Supplementary Fig. 7**) and the chamber was filled with 1 ×PBS before being completely sealed off with vacuum grease.

### Collagen Network Crosslinking

After polymerization of the collagen network as described above, approximately 100 μ*l* of 4 % *v/v* glutaraldehyde (GA) in deionized water, was pipetted on top of the collagen and the sample was placed back into the incubator for ~ 2 *h*. After incubation, the collagen was thoroughly rinsed with 1 ×PBS and the sample chamber was assembled and mounted on the PFM.

### Cell Culture

HeLa cells were kindly gifted to us by Prof. Aaron Baker (Biomedical Engineering Department, The University of Texas at Austin). HeLa cells were adapted to and cultured in CO_2_ independent medium (Gibco™, 18045088), which allows for cell culture under atmospheric conditions. The medium was supplemented with 10 % bovine calf serum (GE, HyClone, SH30072), 4 *mM* L-glutamine, 100 *IU ml^−1^* penicillin and 100 *μg ml*^−1^ streptomycin at 37° *C*. Cells were passaged every three days. To detach cells from the culture flask, the disassociation agent TrypLE™ Express Enzyme 1 × (ThermoFisher Scientific, 12605036) was used.

### Cell Seeding in Collagen Matrix – Polymerization Protocol II

To produce collagen networks with seeded HeLa cells, the collagen network protocol as described above had to be altered in order to incorporate the cell culture medium with a buffer system independent of CO_2_ control as the sample is subject to atmospheric conditions while mounted on the PFM. Similarly to the above, acid-soluble rat-tail tendon collagen (Collagen I, rat tail, 354236, Corning^®^, NY, USA) and bovine-dermis collagen (Collagen I, bovine, 354231, Corning^®^, NY, USA) were mixed at relative concentrations of 1:2 on ice. The mixture was then diluted with 10 % (*vol/vol*) 10 ×PBS and the pH neutralized to pH = 7.2 − 7.4 with 1 *M* NaOH and kept on ice. HeLa cells in culture at 80 – 90 % confluency were detached from the culture flask using the disassociation agent TrypLE™ Express Enzyme 1 ×(ThermoFisher Scientific, 12605036) and spun down at 125 ×*g* for 10 *min*. The cells were then resuspended in serum-free CO_2_ independent medium (Gibco™, 18045088) and supplemented with 4 *mM* L-glutamine at a concentration between (40 – 80) ×10^4^ cells *ml*^−1^. Finally, the medium containing the cells was added to the neutralized collagen solution on ice at a relative concentration of 20 % (vol/vol) to yield a final cell concentration between (7 – 15) ×10^4^ cells ml^−1^, and collagen concentration of 2.4 *mg ml*^-1^. The solution was carefully mixed, pipetted into the preassembled sample chamber and placed in a humidity controlled 37° *C* incubator for 1 h. The network was then topped with ~ 100 μ*l* of serum-free CO_2_ independent cell media and allowed to incubate overnight.

### Calculating the angle of fibril axis

Having determined all locations belonging to the fibril axis, we assign an orientation angle value to each individual location, termed filament location in the following. Choosing one filament location after another as the center, we sweep a full circle of given radius *r* (here: *r* = 4 *pixels*) in discrete angle steps of *π*/100. For every step we calculate an alignment score corresponding to the number of filament locations that are aligned with the straight-line segment connecting −*r* and *r*. The alignment score can take a value between 0 and 2*r* pixels. We then find the angle corresponding to the maximum by Gaussian fitting, which corresponds to the local fibril angle at this location. If a filament location is not connected to any others, therefore alone standing, it is discarded.

### Calculation of Collagen Fibril Fluctuation, Background Correction and Thresholding

While performing Activity Microscopy measurements a small part of the signal can be attributed to background noise, which arises primarily from four different sources: electronic noise in the amplifier and stage control, laser power fluctuations, mechanical instabilities in the PFM setup, as well as background contributions by parts of the network out of focus. Those noise sources could be characterized individually, but it is simpler to estimate the sum background signal by positioning the laser focus inside of network pores in a collagen sample, far away from fibrils. We determined that on average the s.d. of the background signal is *σ̄*_*background,x*_ = 12.56 ± 0.39 *mV* and σ̄_*background,y*_ = 12.33 ± 0.92 *mV*. Assuming all signals to be independent random variables we can subtract the variance of the background signal σ̄^2^_*background,x*_ from the variance of the signal measured in volts *σ*^2^ *S_x_*(*x, y*) to get a fibril’s true motion:

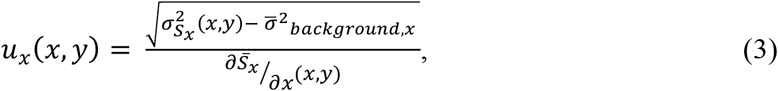

where *∂S̄_x_/∂x* (*x,y*) is the local detector sensitivity. An analogous expression is valid for the *y*-signal. In addition, we chose the smallest acceptable detector sensitivity as 20 *mV pixel^−1^* for pixels with side lengths of 100 *nm*. Any locations with local sensitivity values below our threshold were deleted. Around all measured scanning areas a two pixel-wide frame was deleted as the spatial derivate for finding fibril locations is ill defined at the edges. For larger scans that required the successive scanning of multiple subsections, a 1 μ*m* overlap between the subsections prevented gaps in the data after deleting pixels at the edges.

### Pore Size Distribution for Hot versus Cold Fibrils

An Activity Microscopy image displaying fibril fluctuations is first divided into equally sized sets of “hot” and “cold” fibrils by finding the median of the fluctuation amplitude distribution. We then perform a pore size analysis partially adapted from Mickel *et al.*^19^, on three Activity Microscopy images, albeit in two dimensions. In brief, we first convert the image into a simple binary dataset, where all pixels that belong to a fibril are assigned an arbitrarily high value, and all pixel belonging to a network pore are assigned the value zero. We then determine for every pore pixel the largest disk to fit into a given network pore without intersecting any pixels belonging to a fibril. Each pore pixel in the image is then assigned the radius of the largest disk that incorporated said pixel as its value.

### Data Analysis, Visualization, and Code Availability

The data were acquired and analyzed using custom software written in Labview (National Instruments, TX, USA) and Igor Pro (Wavemetrics, OR, USA) and is available from the corresponding author upon request.

### Data Availability

The data that support the findings of this study are available from the corresponding author upon reasonable request.

## Acknowledgements

We gratefully acknowledge the support of NSF research awards DMR-1310559 and DMR-1710646. This work was partially supported by a Junior Fellow award from the Simons Foundation to T.F.B. We thank Suzanne Jacobs for kindly reading of the manuscript and providing valuable feedback.

## Author Contributions

E.N.L., T.F.B., and E.-L.F. conceived the experiments. E.N.L. performed the experiments, developed analysis software and analyzed data. T.F.B. and E.N.L. wrote software to control the instrument. T.F.B. wrote the software to calculate the angle of a fibril’s axis. E.N.L., T.F.B, and E.-L.F. wrote the manuscript.

